# A synthetic potassium channel reduces oxidative stress via cellular adaptronics

**DOI:** 10.1101/2024.11.18.624138

**Authors:** Alberto Russo, Andrea Saponaro, Simone Trini, Tyler S. Nelson, Heather N. Allen, Rebecca Oddone, Chiara Villa, Yvan Torrente, Rajesh Khanna, Alessandro Porro, Gerhard Thiel, Anna Moroni

## Abstract

Aerobic metabolism is crucial for human life but reactive oxygen species (ROS) byproducts cause cellular toxicity. Although antioxidant defenses usually maintain ROS levels within a safe range, ROS production can exceed the buffering capacity of cells, causing oxidative stress and disease. Inspired by the principle of adaptronics, we created a synthetic potassium channel that senses cellular ROS levels and mitigates oxidative stress by modulating membrane potential. Engineered from TASK1 channel, ROSTASK1 is sensitive to supraphysiological ROS levels, imposing restorative membrane potential changes on cells or organelles under oxidative stress. We also engineered a blue-light sensitive ROSTASK1 to achieve optogenetic control. In proof-of-concept experiments, mitochondrially-delivered ROSTASK1 rescued ROS overproduction in myoblasts from a Leigh syndrome patient and ROSTASK1 abolished chronic pain-like behavior in mouse models of inflammation and nerve injury. Thus, by functioning as both a sensor and modulator of ROS levels, ROSTASK1 provides a self-healing system during oxidative stress.

## Introduction

Reactive oxygen species (ROS) are a byproduct of cellular metabolism, predominantly mitochondrial metabolism, in aerobic organisms. They are also produced in smaller amounts by enzymes such as NADPH oxidase, ionizing and UV radiation, and the metabolism of a wide range of drugs (Juan et al, 2021). Although low concentrations of ROS serve crucial signaling roles in cell differentiation and survival, levels that exceed antioxidant defense systems result in oxidative stress, which can be toxic to cells. Indeed, due to their reactivity with nucleic acids, lipids, and proteins, ROS are associated with cancer, neurodegeneration, and inflammation (Teleanu et al, 2022).

The molecular targets of oxidative stress include proteins involved in the maintenance of essential ion gradients. For example, ROS readily inactivate the Na^+^/K^+^ pump (Andreoli et al, 1993) and can open or inhibit different ion channels, depending on their identity. In addition to direct damage, channel modulation by ROS can have secondary effects that further modify oxidative stress. We hypothesized that engineered ROS-activated channels may offer a novel strategy to sense excessive levels of ROS within cells and mitigate their harmful effects. We focused on K^+^ channels, which are pivotal for controlling the resting membrane potential of cells and subcellular organelles, and therefore key cellular functions.

One example of the beneficial effect of K^+^ channels is the role they play in mitochondria, which produce about 90% of the total amount of cellular ROS due to electrons leaking from the transport chain during hyperpolarization. In physiological conditions, electrons are immediately buffered by enzymes (e.g., SOD) and antioxidants (e.g., vitamin C) (Birben et al, 2012). But when ROS production passes the physiological buffering capacity of a cell, oxidative stress can lead to apoptosis. Mitochondrial K^+^ channels are important in this scenario because they limit ROS overproduction by depolarizing the inner mitochondrial membrane (IMM) (Malinska et al, 2010; Checchetto et al, 2021). We postulated that exogenous ROS-activated K^+^ channels in the IMM could prevent apoptosis during oxidative stress by both detecting elevated ROS levels and depolarizing mitochondria, thus restoring physiological ROS levels.

Another potential role for K^+^ channels in counteracting oxidative stress is during neuropathic pain, a chronic condition caused by lesion or aberrant function of the somatosensory nervous system. Neuroinflammation and oxidative dysfunction contribute to the induction and maintenance of neuropathic pain by increasing the excitability of nociceptors (Ellis et al, 2013; Ji et al, 2018). For instance, ROS can activate neurons that express Ca^2+^-permeable TRPA1 channels, leading to overexcitation (Andersson et al, 2008). In this scenario, opening of ROS-activated K^+^ channels on the plasma membrane of injured neurons would hyperpolarize these cells and inhibit their firing. However, the lack of specific K^+^ channel openers has resulted in significant side effects during studies of candidate pharmaceuticals. The expression of ROS-activated K^+^ channels would be highly advantageous in this context because they would open only when ROS levels are anomalously high due to neuronal damage.

To develop a targeted therapy in which K^+^ channels remain inactive under normal circumstances and become active in pathological conditions to suppress ROS production, we sought to develop a K^+^ channel that opens in the presence of elevated ROS levels. ROS-activated K^+^ channels exist in the voltage-gated K^+^ channel family (Sahoo et al, 2014) and two-pore domain K^+^ channel (K2P) family. TASK1 is one example from the K2P family, but evidence for its modulation by ROS is anecdotal and conflicting (Lee et al, 2006; Kim et al, 2007; Park et al, 2009; Papreck et al, 2012; Turner et al, 2013), and its basal activity would need to be silenced for this application. However, several other factors make TASK1 an ideal candidate for a protein engineering project. Unlike other K2P family members, which are gated only at the selectivity filter, TASK1 possesses a cytosolic X-gate that mechanically adjusts the channel’s open probability. Furthermore, the high-resolution structure of TASK1 (Rödström et al, 2020) offers a solid platform for rational protein engineering, and its small size (45 kDa) permits packaging into AAV viruses for *in vivo* experiments.

In this study, we engineered two TASK1-derived channels, ROSTASK1 and OptoTASK1, that are closed at rest and gated open by elevated ROS levels. These channels are activated by singlet oxygen molecules (^1^O_2_), which irreversibly oxidize two methionine residues in the cytosolic X-gate. They open in response to elevated cellular ROS levels or to light in the presence of a photosensitizer agent. OptoTASK1 incorporates the genetically-encoded photosensitizer miniSOG to optimise the channel for mammalian optogenetics. ROSTASK1 becomes active during oxidative stress and rescues ROS overproduction in a model of mitochondrial dysfunction. Furthermore, ROSTASK1 expression in dorsal root ganglia mitigates mechanical hypersensitivity three models of chronic pain. Our results highlight the considerable potential and diverse applications of ROSTASK1 for ROS-induced ROS control (RIRC) to treat conditions where oxidative stress is an issue.

## Results

### TASK1 is modulated by ^1^O_2_

To confirm earlier observations of TASK1 modulation by ROS, we expressed the channel in HEK293T cells and generated intracellular ROS using UV-A light (Bellono et al, 2013). Ion currents were recorded in the whole-cell patch clamp configuration from control cells and cells transfected with TASK1 (Figure 1A,B). Under safe light, transfected cells showed outwardly rectifying currents in response to voltage steps and a more negative reversal potential (E_rev_= −57 ± 4 mV) than untransfected (UT) cells, consistent with expression of TASK1 channels (Figure 1C,D). Blue light (470 nm) did not affect the amplitude or E_rev_ of TASK1 currents whereas UV-A light (395 nm) increased current amplitudes and shifted E_rev_ further to the left (−67 ± 1.9 mV) in transfected cells (Figure 1C). The effect of UV-A slowly increased over time, reaching an approximately three-fold increase after 10 minutes (half activation time t_1/2_ = 336 ± 20 s) (Figure 1E and Table S1). The antioxidant N-Acetylcysteine (NAC; 5 mM) suppressed this UV-A-induced increase in current, confirming TASK1 modulation by ROS (Figure 1F).

**Figure 1.**
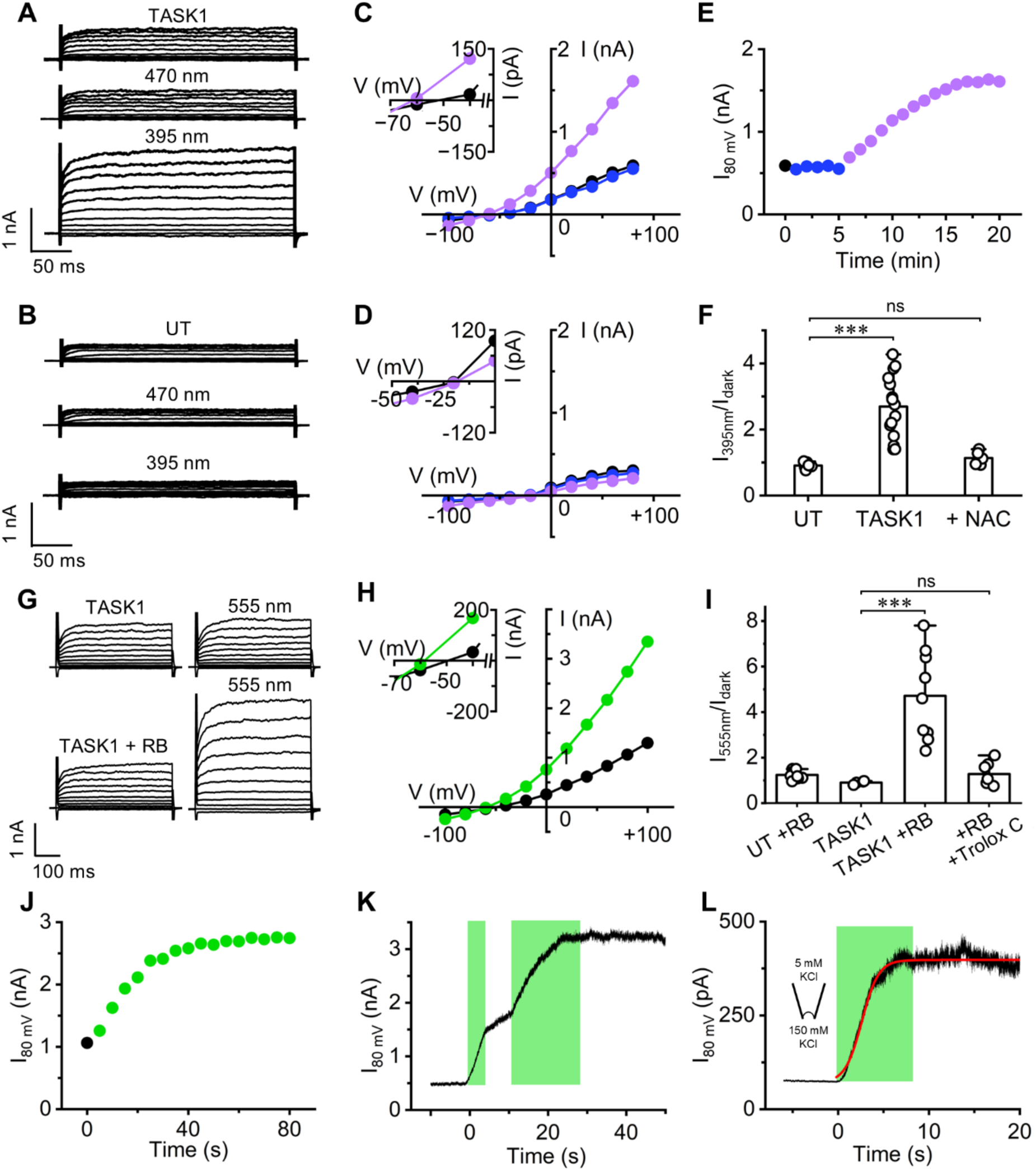
TASK1 activation by ROS. **A, B**: Representative whole-cell recording from HEK293T cells transfected with mTASK1 wt (A) and untransfected cells (UT) (B). Currents were recorded at the end of rundown in safe light condition (top trace), after 5 minutes exposure to 470 nm light (middle trace) and after 15 min exposure to 395 nm light (bottom trace). Light intensity was 55 mW cm^−2^ for both wavelengths. **C, D**: IV relationship from measurement in A (mTASK1 wt) and B (untransfected) in safe light (black), after 5min of 470 nm light (blue) and after 15 min of 395nm light (violet). Inset: zoomed view highlighting E_rev_ for TASK1 (−55 mV in safe light; −67 mV in 395nm) and UT (−20 mV in safe light; −20 in 395nm). Similar results were obtained in n = 4 cells for TASK1 and n=6 for UT. **E**: Time course of current amplitudes measured at 80 mV from cell shown in A before (black circles) and during light exposure (symbols coloured as in panel C). Number of experiments and t_1/2_ values ± SEM are reported in table S1. **F**: Fold change in current amplitude measured at 80 mV after 10 minutes of 395 nm light exposure. Cells expressing TASK1 were measured in control solution (TASK1) or in presence of 5 mM NAC in the extracellular solution (+NAC). n≥ 5 for all conditions. Data are shown as mean ± SEM. One-way Anova with Fisher’s LSD test (***p<0.001; ns, p>0.05). **G:** Representative whole-cell recording of mTASK1 wt in control solution and in presence of 100 nM RB in the extracellular solution before (left) and after 1 min exposure to 555 nm light at 55 mW/cm^2^ (right). **H**: IV relationship from measurement of TASK1 + RB shown in G in safe light (black) and in 555 nm light (green). Inset: zoomed view of X-axis highlighting E_rev_ (−50 mV in safe light; −59 mV in 555 nm). **I**: Fold change in current amplitude measured at 80 mV after 1 minute of 555 nm light treatment for mTASK1 wt in presence of either 100 nM RB (n = 9) or both 100 nM RB and 5 mM TROLOX-C (n= 6) in the extracellular solution. UT cells measured in presence of 100nM RB (n = 9) and mTASK1-expressing cells in control solution (n = 4) are shown as control. Data are presented as mean ± SEM. One-way Anova with Fisher’s LSD test (***p<0.001; ns, p>0.05). **J:** Time course of current amplitudes measured at 80 mV from the cell shown in G before (black symbol) and during exposure to 555 nm light (green symbols). Number of experiments and t_1/2_ values ± SEM are reported in table S1. **K, L**: Representative gap-free recordings measured at +80 mV in whole cell (K) and in inside-out configurations (L) from cells expressing TASK1 in presence of 100 nM RB in the extracellular solution. The green bars indicate the time span during which 555 nm light was delivered. Similar results were obtained in n ≥ 3 cells. Current data were fitted with a dose-response function (red line in panel L).

UV-A radiation interacts with chromogenic groups and photosensitizers in cells, producing the non-radical ROS ^1^O_2_, which is rapidly metabolized to free radicals such as superoxide anion radical (•O2^−^) and hydroxyl radical (•OH) (Hanson and Simon, 1998). We investigated whether these free radicals affect TASK1 conductance by treating cells with the oxidizing agent hydrogen peroxide (H_2_O_2_), which enters cells and rapidly converts into •O2^−^ and •OH. After 15 minutes, 10 mM H_2_O_2_ had no impact on TASK1 currents (Figure S1), ruling out a major role for H_2_O_2_ and its by-products in channel modulation.

We subsequently examined the effect of Rose Bengal (RB), a photosensitizer known to produce high levels of ^1^O_2_ when exposed to green light (550 nm) (Kochevar and Redmond, 2000; Gao et al, 2014). Upon green light exposure, 100 nM RB stimulated strong outward currents and a marked negative shift of E_rev_ in cells expressing TASK1 (Figure 1G,H). This effect was blocked by 5 mM Trolox C, a specific ^1^O_2_ quencher (Figure 1I), suggesting that ¹O₂ is the ROS that regulates TASK1 activity. The current induced by RB in the presence of green light increased > 20 times faster than that induced by UV-A (Figure 1J, Table S1). In addition, the degree of current activation was dependent on the duration of light exposure (Figure 1K). Experiments using inside-out macro-patches revealed that intracellular exposure to RB in the presence of green light increased the speed of activation (t_1/2_ ~ 3 s) (Figure 1L), and moreover, that RB-induced modulation does not depend on intracellular components. Notably, the finding that RB-induced currents are stable after green light is turned off indicates a long-lasting effect (Figure 1K,L). These findings suggest that the ROS ^1^O_2_ rapidly and persistently activates TASK1 channels expressed in HEK293T cells.

### ROSTASK1 is activated by ROS

A mammalian ROS-activated K^+^ channel could be a useful tool in research and therapy if the channel was otherwise closed. Because TASK1 is open at rest, we sought to engineer a cytosolic gate to prevent basal activity. The resulting construct, ROSTASK1, is a truncated version of mouse TASK1 incorporating eGFP at the C-terminus for cellular localization (Figure 2A). The construct was truncated at E259 to remove known regulatory phosphorylation domains in the C-terminus (Zuzarte et al, 2009; Seyler et al, 2012). Point mutations Q126M and T248F were introduced to reinforce the hydrophobic interactions between the transmembrane helices in the region where the M4 helices cross (X-gate) (Figure 2A, Figure S2A), thus removing residual channel activity (Rödström et al, 2020). Specifically, the Q126M mutation on M2 reinforced the hydrophobic interaction with L244 on the same monomer (M4) and M247 on the opposite monomer (M4’) (Figure S2B). Furthermore, T248F on M4 reinforced the hydrophobic interaction between itself and the same residue on M4’ to strengthen the X-gate and fully close the channel (Figure 2A).

**Figure 2.**
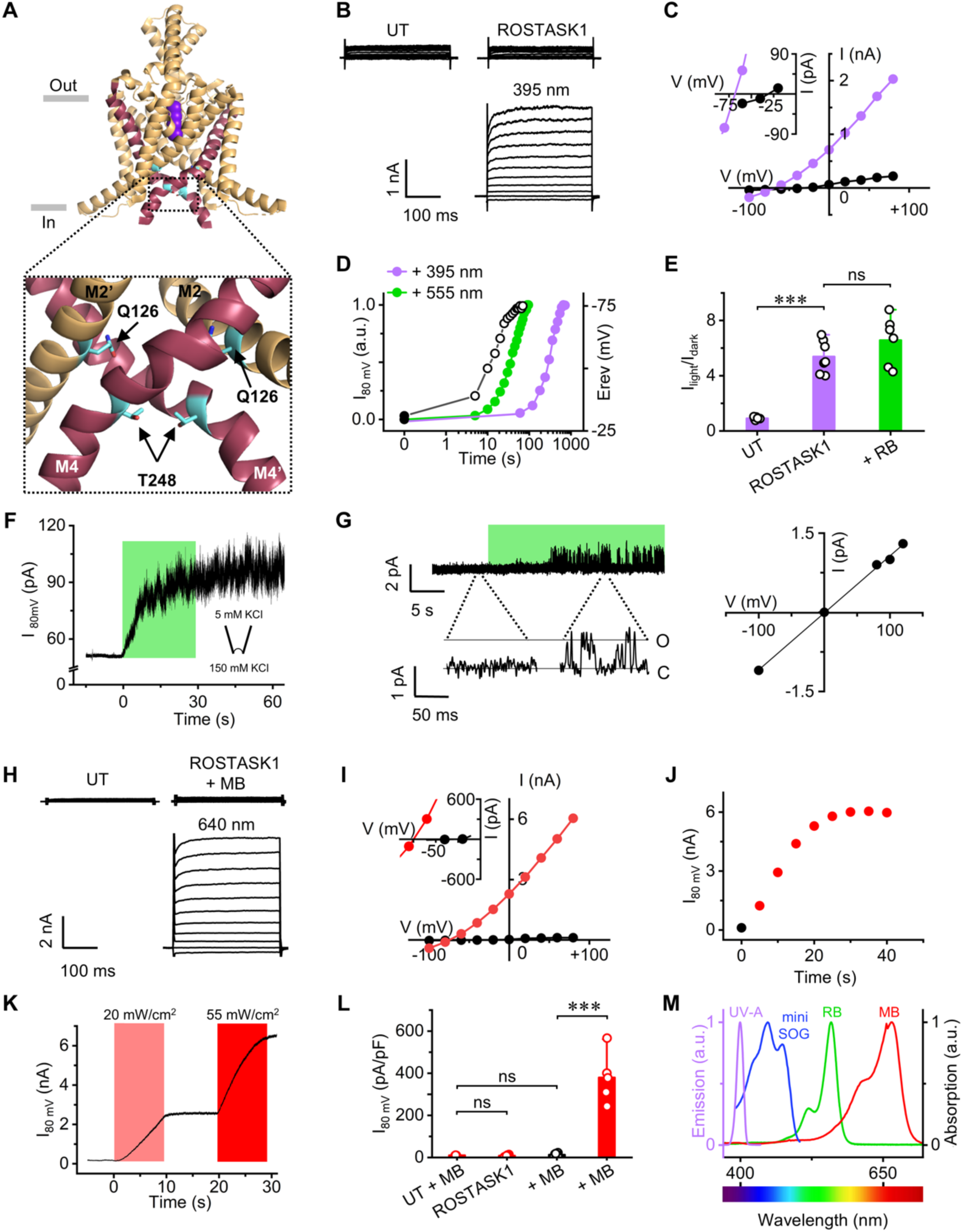
Engineering of ROSTASK1 and MB shifts the dynamic range of activation towards far red light. **A**: Top, front view of the truncated (Met1-Glu259) human TASK1 structure (PDB: 6rv2). M4 helices are shown in red to highlight the X-gate. Side chains of Q126 and T248 residues are in light blue. Grey lines indicate plasma membrane boundaries. Boxed, enlarged view of the region surrounding the X-gate. For clarity, only M2 and M4 helices from both chains are shown. **B**: Representative whole-cell current traces of ROSTASK1 in safe light (top) and after 15 min exposure to 395 nm light (bottom). An untransfected cell (UT) measured in safe light is shown for comparison. **C**: IV relationship from ROSTASK1 current shown in B before (black) and after 395nm light exposure (violet). Number of experiments: 7. Inset: zoomed view highlighting E_rev_ (−30 mV in safe light; −68 mV in 395nm). **D**: Representative time course of normalized currents measured at 80 mV for ROSTASK1 exposed to 395 nm light (same cell shown in B) or 555 nm light (55 mW/cm^2^ for both wavelengths). Cell exposed to 555 nm light was measured in presence of 100 nM RB in the extracellular solution. Black empty symbols show time course of E_rev_ shift measured from the same cell exposed to 555 nm light. Full black symbol at t0 shows current in safe light condition. Number of experiments and t_1/2_ values ± SEM are reported in table S1. **E**: Fold change in current amplitude measured at 80 mV in cells expressing ROSTASK1 after 10 minutes of 395 nm light (ROSTASK1, n=7) or after 1 minute of 555 nm light exposure (+RB, n=6). UT cells exposed to 395 nm light are shown as control (n= 6). Data are presented as mean ± SEM. One-way Anova with Fisher’s LSD test (***p<0.001; ns, p>0.05). **F**: Representative gap-free recording measured at 80 mV in inside-out configuration from a cell expressing ROSTASK1 with 100 nM RB. The green bar indicates the time span 555 nm light was delivered. **G**: Left, single channel recordings measured at 120 mV in cell-attached configuration from a cell expressing ROSTASK1 with 100 nM RB. The green bar represents the time span during which light was delivered. The magnifications show current in safe light (no channel activity observed) and after 555 nm light treatment. Right, IV plot of single channel current amplitudes measured at five different voltages. Data fit to linear equation provided a conductance of ~10 pS (similar to that reported for TASK1 wt). **H**: Representative whole-cell current traces of ROSTASK1 in safe light (top) and after 1 min exposure to 640 nm light (bottom) in presence of 5 µM MB in the extracellular solution. An untransfected cell (UT) measured in safe light is shown for comparison. **I**: IV relationship from the ROSTASK1 trace shown in H. Similar results were obtained in n ≥ 4 cells. Inset: zoomed view highlighting E_rev_ (−27 mV in safe light; −75 mV in 640 nm). **J**: Time course of current amplitudes measured at 80 mV from cell shown in H before (black circle) and during 640 nm light exposure (red). Light intensity: 55 mW/cm^2^. Number of experiments and t_1/2_ values ± SEM are reported in table S1. **K**: Whole-cell gap-free recording measured at 80 mV from a cell expressing TASK1 with 5 µM MB. The red bars indicate the time span during which 20 and 55 mW cm^−2^ 640 nm lights were delivered. **L**: Current densities measured at 80 mV in safe light (black) and after 1 min light 640 nm light exposure at 55 mW cm^−2^ from untransfected cells + 5 µM MB (UT + MB, n = 3), ROSTASK1 in control solution (n = 4), ROSTASK1 + 5 µM MB (MB, n ≥ 5). One-way Anova with Fisher’s LSD test (***p<0.001; ns, p>0.05). **M**: Normalized absorbance of miniSOG, RB and MB and plotted along with the emission spectra of UV-A light generated by SPECTRA X light engine device (Lumencor).

Basal currents recorded from ROSTASK1 were indistinguishable from those of untransfected cells in terms of amplitude and E_rev_ (Figure 2 B), verifying that the channel is closed at rest. However, UV-A light activated large currents in ROSTASK1-transfected cells, confirming its expression at the plasma membrane and, importantly, retention of the ROS activation mechanism. Current activation was accompanied by a shift of E_rev_ to −69 mV (Figure 2C), close to the Nernst potential for K^+^ (E_K_), and the t_1/2_ for UV-A (431 ± 88 s) was similar to that of the wild type channel (Figure 2D). The combination of green light and RB activated the channel about 20 times faster (t_1/2_ = 25 ± 5.4 s) than UV-A, reaching similar activation levels (Figure 2D,E). Interestingly, the change in E_rev_ was evident after a few seconds of light and preceded any measurable increase in current (Figure 2D). This suggests that E_rev_ is a more sensitive indicator for channel activation. As observed for TASK1, the effect of light was faster in inside-out patches and persisted after the light was turned off (Figure 2F). In addition, the fold change in current through ROSTASK1 was similar for activation by UV-A and green light plus RB, as was the case for TASK1 (Figure 2E). Single channel recordings in cell-attached patches with 100 mM KCl in the pipette solution revealed that channels were closed in safe light and opened after about 10 s of green light plus RB stimulation (Figure 2G). Channel conductance was 10 pS, as expected for TASK1 channels in symmetrical 100 mM KCl solutions (Coeli et al, 2000) (Figure 2G).

Methylene blue (MB), also known as methylthioninium chloride, is a hydrophilic phenothiazine derivative and a known antimicrobial agent (Bueno-Silva et al, 2024). It is a photosensitizer with a light absorption peak in the red spectrum (660 nm) that generates ROS (Chen et al, 2021). MB (5 µM) robustly and rapidly activated ROSTASK1 (t_1/2_ = 11 s) (Figure 2H) with low light requirement (20 mW/cm^2^) (Figure 2K), shifting E_rev_ towards E_K_ (Figure 2H,I), whereas red light or MB alone had no effect (Figure 2L). MB is therefore a tool that expands the wavelengths for ROSTASK1 activation towards far red spectrum, which is highly desirable for light penetration into tissue. Because blue wavelengths could also be included by incorporation of the genetically-encoded photosensitizer protein miniSOG, as described below, ROSTASK1 has the potential to be activated by the entire visible spectrum of light (Figure 2M). Together, these experiments reveal that ROSTASK1 is a TASK1 channel that is closed at rest and activated by an increase in ROS.

### OptoTASK1 is a blue light-activated channel

ROSTASK1 is a bona fide optogenetic tool that can be activated by green and red light when a photocatalyst is present. To create a genetically-encoded optogenetic tool that does not require a photosensitizer, we replaced the C-terminal eGFP with the photosensitizer miniSOG (Figure 3A). MiniSOG is a small (14 kDa) fluorescent flavoprotein, engineered to generate ^1^O_2_ when illuminated by blue light (Shu et al, 2011). The putative structure of this new channel, named OptoTASK1, incorporates two miniSOG proteins in a functional dimer (Figure 3A). ROSTASK1- and OptoTASK1-transfected cells show similarly small endogenous currents in safe light, but illumination with blue light activates large currents only through OptoTASK1, shifting E_rev_ towards E_K_ (Figure 3B,C). This light-activation of OptoTASK1 persisted for at least four hours after light treatment (15 minutes at 55mW/cm^2^) and was no longer measurable after eight hours (Figure 3D). This timing corresponds to the estimated half-life of a K^+^ channel at the plasma membrane of HEK cells (Colley et al, 2007), suggesting that ROS-induced oxidation is irreversible and its effect is measurable while the channel is present at the plasma membrane.

**Figure 3.**
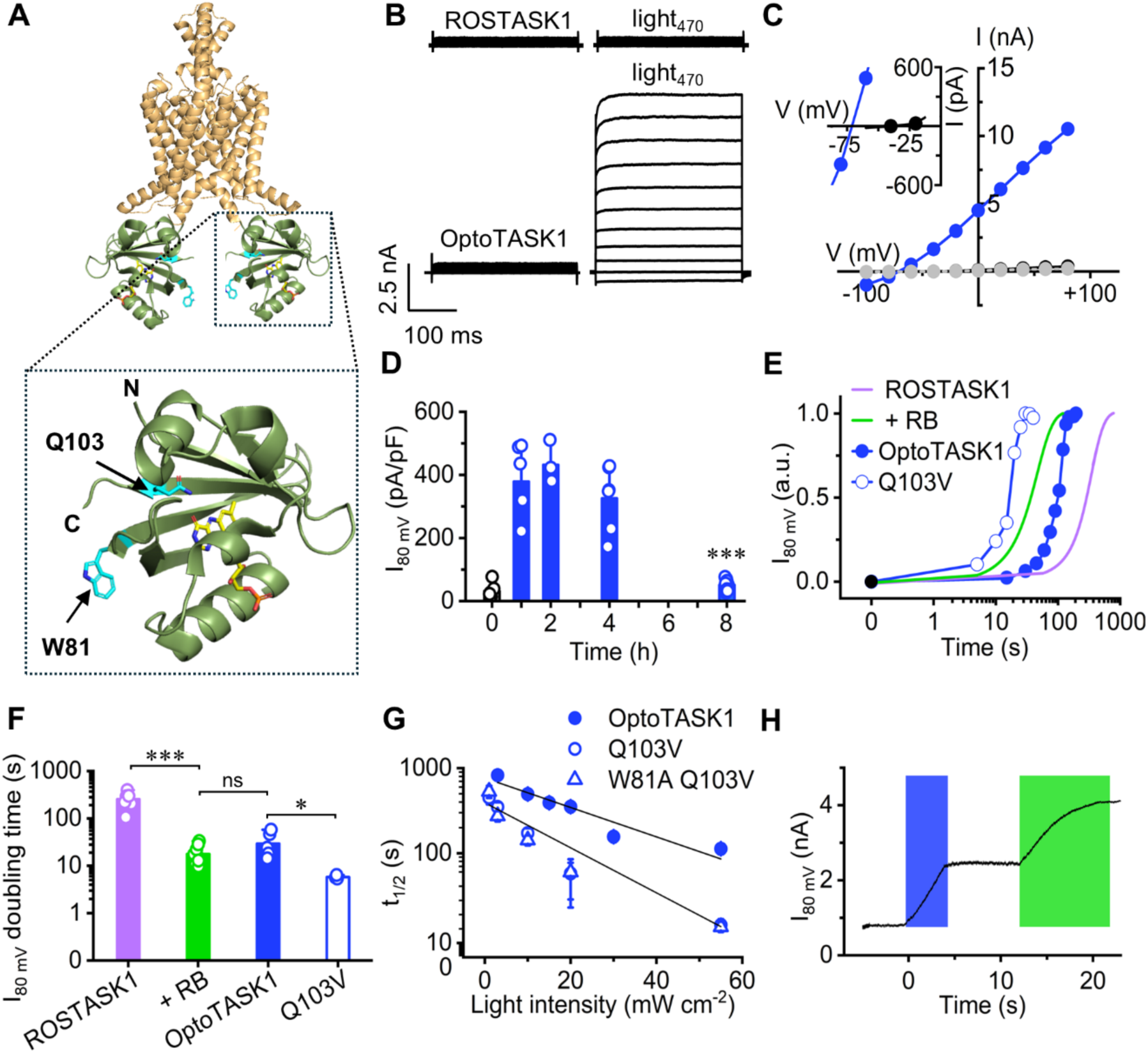
Engineering of OptoTASK1. **A**: Hypothetical structure representing OptoTASK1 generated by fusing C-term of the truncated mTASK1 channel (Met1-Glu259) (ochre) with the N-term of the photosensitizer miniSOG (green). Boxed, miniSOG (PDB: 6gpu) with the FMN chromophore shown in yellow. W81 and Q103 residues are shown as sticks in light blue. **B:** Representative current traces of ROSTASK1 and OPTOTASK1 in safe light (left) and after 150 s illumination with blue light (470 nm, 55 mW cm^−2^) (n = 8). **C**: IV relationships of OptoTASK1 traces shown in B in safe light (black) and in 470 nm light (blue). ROSTASK1 trace in safe light is shown in grey. Inset: magnified portion showing E_rev_ (−40 mV in safe light; −71 mV in blue light). **D**: Time course of OptoTASK1 current density measured at 80 mV. Cells were exposed to 470 nm light for 15 minutes and measured in safe light after 1h (n = 6), 2h (n = 4), 4h (n =6) and 8h (n = 9) hours (blue bars). Black bar at t0 shows current density before light treatment (n=6). Data are presented as mean ± SEM. One-way Anova with Fisher’s LSD test (***p<0.001; ns, p>0.05). **E**: Representative time course of normalized current measured at +80 mV for ROSTASK1 with 395 nm light (violet, replotted from Fig 2D), ROSTASK1 + 100 nM RB with 555 nm light (green, replotted from Fig 2D), OptoTASK1 with 470 nm light (full blue circles, n = 8) and OptoTASK Q103V with 470 nm light (open blue circles, n=3). Number of experiments and t_1/2_ values ± SEM are reported in table S1. **F**: Doubling time of the current in cells expressing ROSTASK1 (n = 6), ROSTASK1 + 100 nM RB (n = 9), OptoTASK1 (n = 7) and OptoTASK1 Q103V (n = 3) treated with 395, 555, 470 nm light, respectively. Data are presented as mean ± SEM. One-way Anova with Fisher’s LSD test (***p<0.001; *p<0.05; ns, p>0.05). **G**: half activation time (t_1/2_) values plotted as a function of blue light intensities (470nm at 1, 3, 10, 15, 20, 30 and 55 mWcm^−2^). t½ values were calculated by fitting rates of activation (see Table S1). Data from OptoTASK1 wt and Q103V mutant were fitted with a linear function (black lines). Number of experiments and t_1/2_ values ± SEM are reported in table S1. **H**: Whole cell gap-free recording of OptoTASK1 with 100 nM RB measured at +80 mV. The blue and green bars indicate the time span during which 470 and 555 nm light were delivered.

Activation by blue light (t_1/2_ = 108 ± 12 s) was four-fold faster than ROSTASK1 activation by UV-A light (Figure 3E and Table S1) but slower than ROSTASK1 activation by green light plus RB. To improve these activation kinetics, we tested two mutations in miniSOG (Q103V and W81A) that are known to increase the rate of ROS production (Westberg et al, 2017). A single Q103V mutation decreased t_1/2_ to 15 ± 4.2 s and reduced the current doubling time to < 10s (Figure 3E,F), while subsequent insertion of W81A to generate the double mutant had no further effect. Although these measurements were obtained at a high light intensity of 55 mW/cm^2^, the activation kinetics for the single and double OptoTASK1 mutants remained relatively fast (t_1/2_ = 60 s) at a low light intensity of 20 mW/cm^2^, due to the considerable light sensitivity of miniSOG (Figure 3G). Importantly, we observed dual activation of OptoTASK1 by either blue light or green light plus RB (Figure 3H), revealing the suitability of our construct for multiplexed optogenetics. Thus, OptoTASK1 is a light-activated K^+^ channel and a valuable optogenetic tool due to its low light requirement, long-lasting activation, and sensitivity to multiple wavelengths of light. These experiments further confirmed that ROSTASK1 channels are irreversibly activated by ^1^O_2_.

### Blue light destabilizes the X-gate in OptoTASK1

ROS typically affects proteins by oxidizing cysteine, methionine, tyrosine, or histidine residues. However, because methionine is preferentially and irreversibly oxidized by ^1^O_2_ (Kim et al, 2013; Rosenfeld, 2023), we considered whether methionine residues in ROSTASK1 and OptoTASK1 might be targets of ROS activation. The addition of an oxygen atom to methionine’s thiol group would generate methionine sulfoxide and a second oxygen atom would further oxidize the sulfoxide to sulfone. Although methionine sulfoxide reductases can repair methionine sulfoxide, conversion to sulfone is irreversible *in vivo*. In TASK1, two methionines on the M3 helix (Met156 and Met159) form a hydrophobic clamp that stabilizes the side chain of Leu241 on the facing M4 helix (Figure 4A). This leucine forms the hinge that gives rise to the X-gate and is therefore critical for gating (Rödström et al, 2020). When mutated into alanine, it promotes channel opening, likely due to weakened hydrophobic interactions and a straightening of the M4 helix. Oxidation of methionine to sulfoxide and then sulfone would lead to a progressive increase in hydrophilicity of the residues, and consequently, destabilization of the hydrophobic clamp that keeps Leu241 in place.

**Figure 4.**
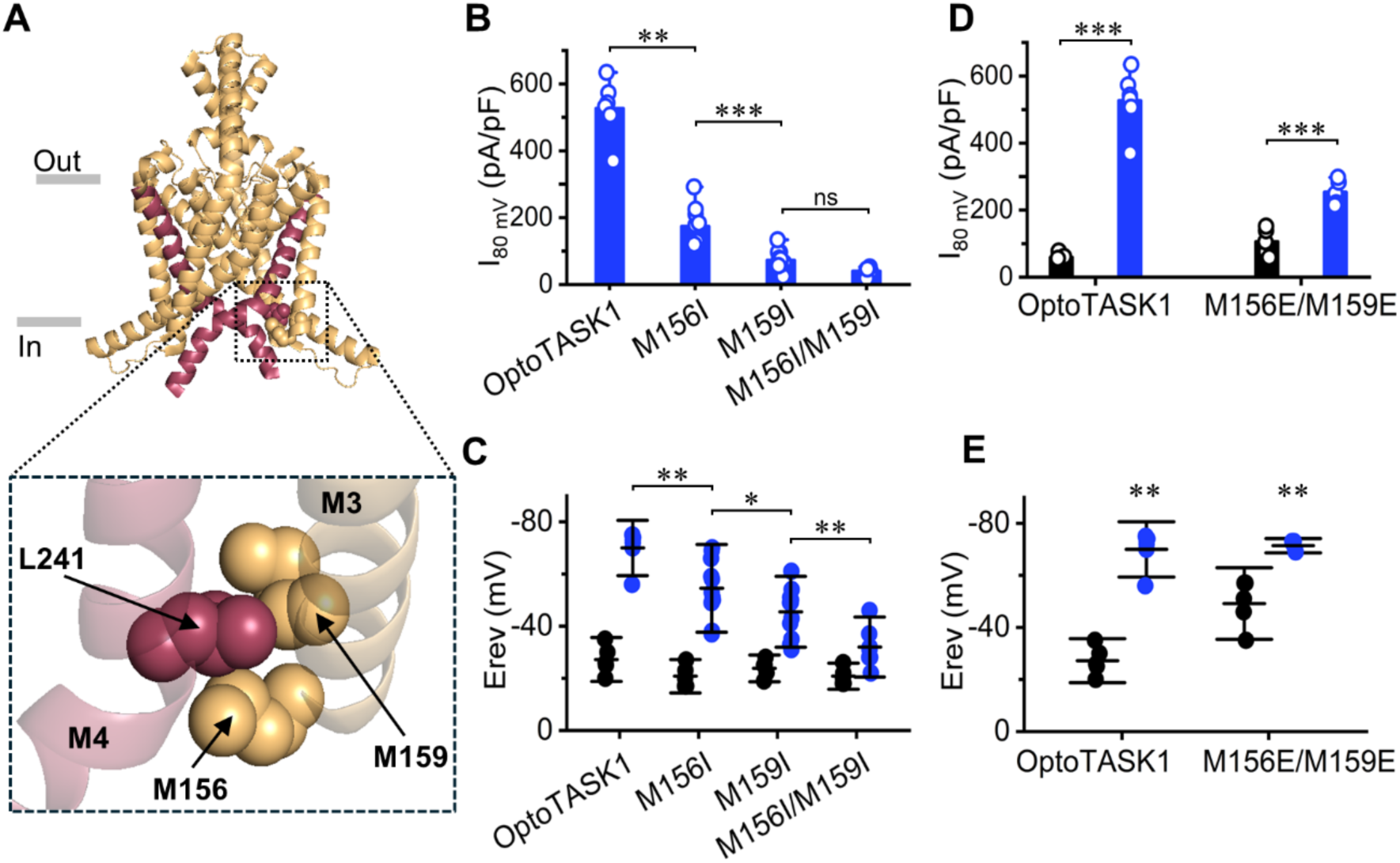
ROS-induced irreversible oxidation of M156 and M159 opens the channel gate. **A**: Top, front view of the structure of the truncated human TASK1 channel (PDB: 6rv2). M4 helices from both chains are shown in red to highlight the X-gate. M156 and M159 on the M3 helix from one chain and L241 on M4 are shown as spheres. Boxed: expanded view of the kink in the M4 helix at the beginning of the X-gate. **B**: Current densities measured at 80 mV in cells expressing OptoTASK1 wt (n=6), M156I (n=9), M159I (n=12) and M156I/M159I (n=7) after treatment with 470 nm light (10 mW/cm^−2^) for 15 minutes. Data are shown as mean ± SEM. One-way Anova with Fisher’s LSD test (***p<0.001; **p<0.01; ns, p>0.05). **C**: E_rev_ measured from the same cells shown in B before (black) and after 470 nm light treatment (blue). Dataset with 470nm light is as in panel B. Number of experiments in dark: n=5 for each group. Data are shown as mean ± SEM. One-way Anova with Fisher’s LSD test (**p<0.01; *p<0.05). **D**: Current densities measured at +80 mV in cells expressing OptoTASK1 wt (n=≥5) and M156E/M159E (n=≥5) in safe light (black) or after treatment with 470 nm light (10 mWcm^−2^) for 15 minutes (blue). OptoTASK1 data in light are replotted from panel B. Data are shown as mean ± SEM. Student’s t-test (***p<0.001). **E:** E_rev_ measured from the same cells shown in D. Data are shown as mean ± SEM. Student’s t-test (**p<0.01).

To test whether oxidation of M156 and M159 underlies ROS-mediated modulation, we substituted one or both methionine residues with isoleucine, which has similar hydrophobicity to methionine but cannot undergo side chain oxidation. Both M156I and M159I in OptoTASK1 drastically reduced blue light-evoked currents and the negative shift in E_rev_. M159I caused the strongest reduction in current density (at 80 mV) and E_rev_ shift, compared to OptoTASK1 (Figure 4B,C). The double mutant was almost insensitive to blue light, suggesting that M156 and M159 are responsible for the majority of OptoTASK1’s sensitivity to ^1^O_2_. Notably, E_rev_ values for the single and double mutants in safe light did not differ from OptoTASK1, indicating that these mutations did not alter channel gating *per se* (Figure 4C).

To confirm our hypothesis that ^1^O_2_ affects gating by oxidizing the thiol groups of M156 and M159 to sulfoxide and sulfone, we replaced these residues with glutamic acids to mimic the negative charges acquired upon oxidation. In contrast to OptoTASK1, the double mutant (M156E/M159E) forms an open channel in safe light, as predicted. However, blue light caused some residual activation and a small change in E_rev_ (Figure 4D,E), indicative of the fact that the single negative charge associated with glutamic acid does not fully mimic the addition of two negative charges upon double oxidation of methionine to sulfone and/or that other oxidizable residues are marginally involved in TASK1 gating besides the two methionine we have mutated. Nevertheless, our data reveal that activation of OptoTASK1 by ^1^O_2_ is primarily due to oxidation of M156 and M159 and thus destabilization of the X-gate.

### mtROSTASK1 limits ROS production by mitochondria

Given that mitochondria produce 90% of cellular ROS, and K^+^ channels can mitigate oxidative stress by regulating mitochondrial membrane potential, we sought to deliver our ROS-activated channels to the IMM to assess their potential impact. We fused six repeats of the mitochondrial-targeting sequence from subunit VIII of human cytochrome c oxidase (Cox-8) to the N-terminal sequence of ROSTASK1 and OptoTASK1. Because TASK1 is inhibited by protons from the redox chain across the IMM, we also mutated the responsible histidine into an asparagine (H98N) (Yuill et al, 2004), significantly reducing the pH sensitivity of the channel (Figure S3 A,B). The resulting constructs, mtROSTASK1 and mtOptoTASK1, colocalized with the voltage-sensitive mitochondrial marker TMRM when expressed in HEK293 cells (Figure S3C). In contrast, ROSTASK1 clearly colocalized with the plasma membrane marker CellMask, ruling out any trafficking defects. Importantly, the bright red signal observed after transfection of these channels (Figure S3C) is indistinguishable from that of untransfected cells (Figure 5A), indicating hyperpolarized and functional mitochondria. Thus, our mitochondrially-targeted ROS-activated K^+^ channels are correctly delivered to the IMM and remain closed with physiological levels of ROS.

**Figure 5.**
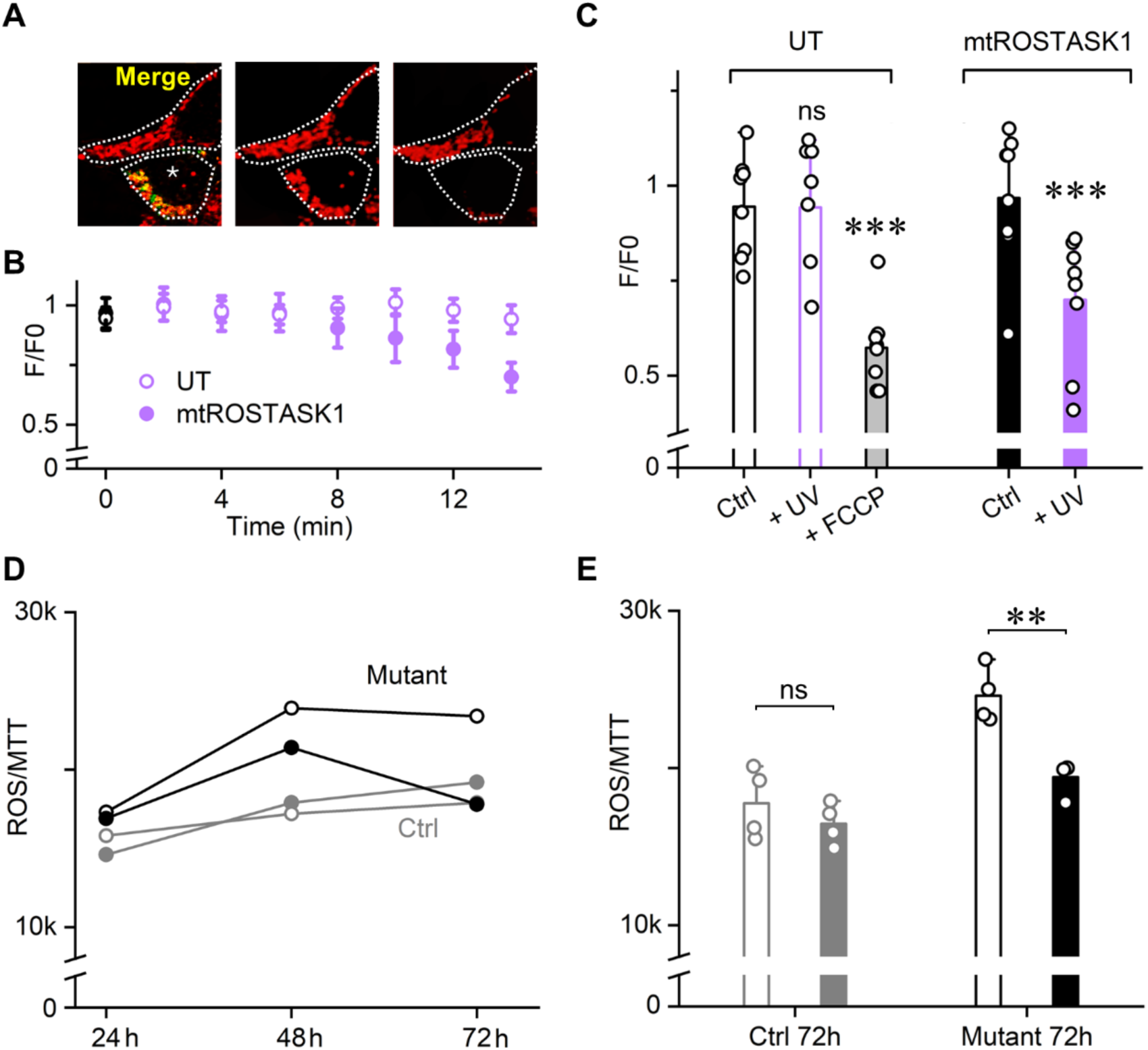
Physiological effects of Mitochondrial ROSTASK1 activation. **A:** mtROSTASK1 activation depolarizes the membrane potential in mitochondria. Left, fluorescent confocal images of an untransfected HEK293T cell and a cell expressing mtROSTASK1 (indicated with a star) showing TMRM red signal in mitochondria merged with EGFP signal of mtROSTASK1. Cells boundaries are marked with a dotted white line. Center: red fluorescence image of the same cells in safe light condition (green signal of mtROSTASK1 was removed for clarity). Right: after 15 minutes of illumination with UV-A light (405 nm at 30 mWcm^−2^). **B:** Time course of normalized TMRM fluorescence in untransfected cells (UT, n=8) and in cells expressing mtROSTASK1 (n=8). Data are shown as mean ± SEM. Black symbol at t0 indicates fluorescence in safe light. **C:** Mean normalized fluorescence intensities of measurements shown in B in safe light (black bars) and after 15 minutes of 405nm light treatment (violet bars). Gray bar: UT cells in presence of 50 uM FCCP in the extracellular solution (which leads to mitochondrial depolarization) measured after 15 min 405 nm light treatment. One-way Anova with Fisher’s LSD test (***p<0.001; **p<0.01; ns, p>0.05). **D**: Total ROS levels in the culture media of healthy (Ctrl, gray) and m.9176T→C (Mutant, black) myoblasts, quantified as the ratio of ROS production over the MTT vitality assay, after 24, 48 and 72 hours after transfection with GFP (empty symbols) or mtROSTASK1 (full symbols). **E:** Mean ROS/MTT levels in the culture media of healthy and m.9176T→C myoblasts at 72 hours after transfection with GFP (gray) or MtROSTASK1 (black). n= 4 for each experimental group. Student’s t-test (**p<0.01; ns, p>0.05).

We next tested whether opening of mtROSTASK1 channels would depolarize the IMM potential by measuring changes in TMRM fluorescence upon channel activation by UV-A light (Figure 5A). In the presence of mtROSTASK1, UV-A treatment led to a decrease in the ratio of TMRM fluorescence in transfected and untransfected cells (Figure 5B), indicating mitochondrial depolarization at a similar rate to that of channel activation by UV-A light (Figure 2D). The degree of depolarization was also comparable to that induced in untransfected cells by the mitochondrial uncoupler FCCP (50 µM) (Figure 5C).

Multiple mitochondrial disorders are associated with oxidative stress and neurological deficits or neurodegeneration, including Leigh syndrome (LS) (Tanaka et al, 1996). To investigate whether mtROSTASK1 activation could mitigate ROS overproduction in LS, we conducted experiments in healthy human skeletal myoblasts and myoblasts from a patient affected by LS with a homoplasmic m.9176T→C mitochondrial mutation, whose muscle tissues exhibited COX-deficient muscle fibers and Complex V dysfunction (Ronchi et al, 2011). Both myoblast lines were transfected with either eGFP or mtROSTASK1, and the ROS H_2_O_2_ measured and normalized to the activity of MTT (a marker of metabolic activity). In all cases, H_2_O_2_ was evident in the culture media, but levels were higher in m.9176T→C myoblasts, indicating that healthy myoblasts can more efficiently control ROS production (Figure 5D,E). Expression of mtROSTASK1 significantly reduced elevated ROS levels in m.9176T→C myoblasts, restoring levels to those observed in healthy controls (Figure 5D,E). Interestingly, there was no obvious effect of mtROSTASK1 in healthy cells, in agreement with our hypothesis that mtROSTASK1 is not activated at physiological levels of ROS levels. These data demonstrate that mtROSTASK1 can mitigate ROS production by depolarising mitochondria, supporting its potential utility in pathological conditions associated with mitochondrial dysfunctions.

### ROSTASK1 alleviates mechanical hypersensitivity in chronic pain states

ROS play a pivotal role in the initiation and maintenance of chronic neuropathic and inflammatory pain (Yowtak et al; Salvemini et al, 2011; Zhang et al, 2023; Xu et al, 2021). We therefore evaluated whether ROSTASK1 could alleviate pain-related behavioral symptoms *in vivo* by intrathecally transfecting dorsal root ganglia (DRG) neurons in adult male and female C57BL/6J mice with either AAV9-CAG-ROSTASK1 or a control AAV9-CAG-GFP vector (Skorput et al, 2022) (Figure S4). Three weeks post-administration, we observed no discernible differences between the animals’ ability to promptly react to a noxious thermal stimulus (52 °C hotplate) or maintain ambulatory function on an accelerating rotarod (Figure 6A, B). To assess the impact of ROSTASK1 on pain sensitivity, we induced persistent inflammation, chronic neuropathic pain, or carried out sham injury in both groups of mice. Persistent inflammation was induced via intraplantar injection of Complete Freund’s Adjuvant (CFA) and chronic neuropathic pain using the spared nerve injury (SNI) paradigm (Nelson et al, 2022).

**Figure 6.**
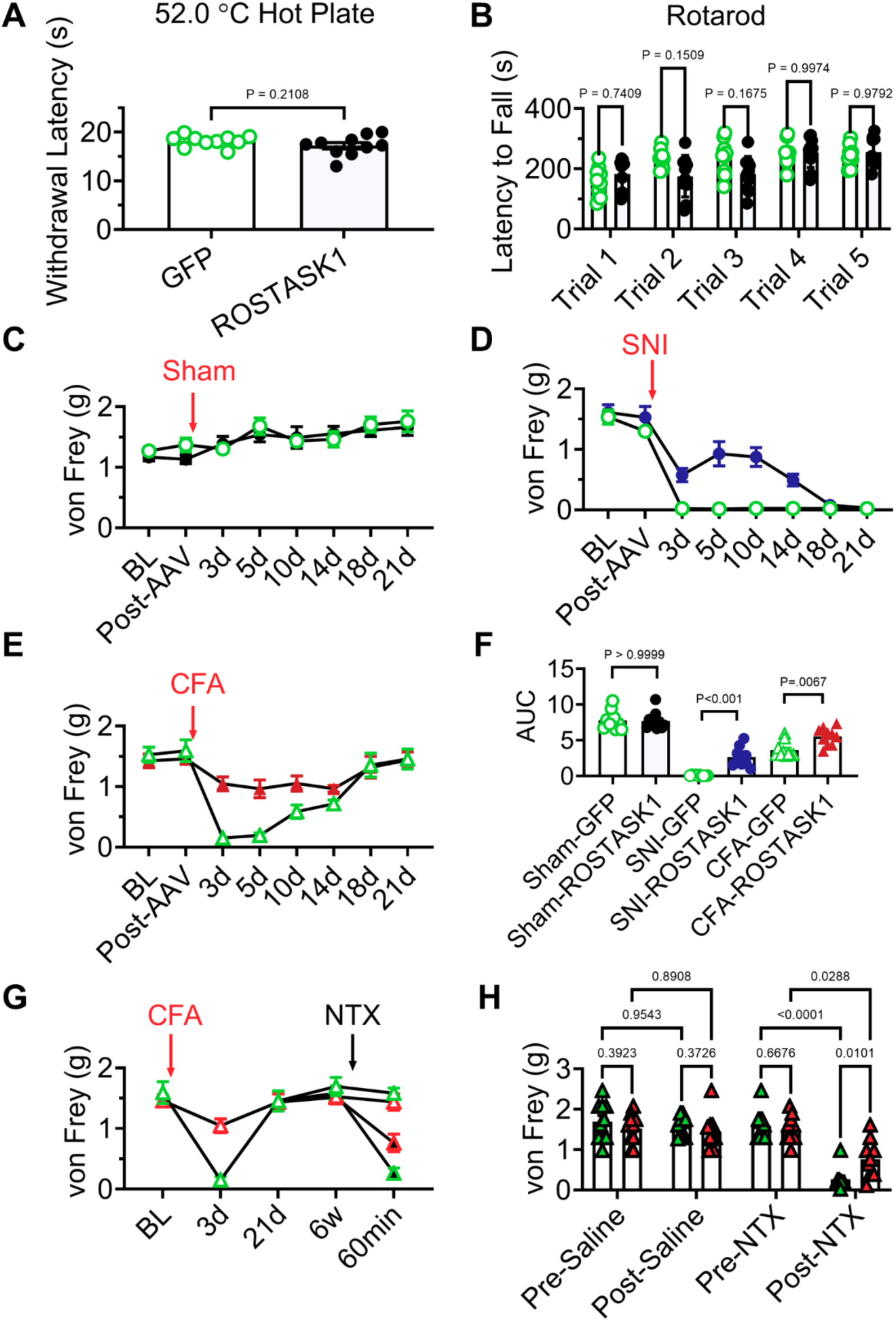
ROSTASK1 alleviates mechanical hypersensitivity in chronic pain states. Behavioral tests for selectively targeting the pain neuraxis performed 21 days after intrathecal injections of AAV9-CAG-GFP (GFP) or AAV9-CAG-ROSTASK1 (ROSTASK1) (see also Figure S4). **A** ROSTASK1 (black) does not affect acute heat sensitivity as assessed by withdrawal latency on a 52.0 °C hotplate. n = 10 mice/group. Data are shown as means ± SEM. Student’s unpaired two-tailed t test. **B:** ROSTASK1 (black) does not affect motor coordination as assessed by latency to fall on an accelerating rotarod. n = 10 mice/group (Two-way RM ANOVA; Virus x Trial; F. (4,72) =4.365; Šidák’s multiple comparisons test). Data are shown as means ± SEM. Dots represent data points from individual animals. **C:** Mechanical sensitivity von-Frey test in GFP mice (green symbols) and ROSTASK1 mice (black symbols) that underwent sham injury (see methods); n =10 mice/group. **D:** Mechanical sensitivity von-Frey test in GFP mice (green symbols) and ROSTASK1 mice (red symbols) that underwent SNI surgery to induce chronic neuropathic pain; n =10 mice/group. **E:** Mechanical sensitivity von-Frey test in GFP mice (green symbols) and ROSTASK1 mice (blue symbols) that received intraplantar injection of CFA to induce persistent inflammation; n =10 mice/group. **F:** Area under curve (AUC) of data shown in panels C-E. Two-way RM ANOVA; Virus x Injury; F. (4,72) =7.523; Šidák’s multiple comparisons test. Data are shown as means ± SEM. **G:** Mechanical sensitivity von-Frey test in GFP (green) and ROSTASK1 mice (red) groups shown in panel E. Naltrexone (NTX, filled symbols) or Saline (empty symbols) was injected 6 weeks after CFA to reinstate the mechanical hypersensitivity. Data are shown as means ± SEM. **H**: Mechanical sensitivity von-Frey test before and after NTX injection in mice groups shown in panel G. Dots represent data points from individual animals. Two-way RM ANOVA; Virus x Drug Time Point; F. (3,54) =3.882; Holm Šidák’s multiple comparisons test.

In sham-treated mice, AAV9-CAG-ROSTASK1 had no effect on mechanical sensitivity, measured using von Frey filaments (Figure 6C). However, AAV9-CAG-ROSTASK1 significantly reduced the development of CFA-induced mechanical allodynia (Figure 6E) and mitigated SNI-induced mechanical allodynia (Figure 6D), compared to AAV9-CAG-GFP mice. The anti-allodynic effect of AAV9-CAG-ROSTASK1 on SNI-induced mechanical hypersensitivity diminished after 14 days post-injury, suggesting a transient role of ROS in the maintenance of long-term neuropathic pain (Figure 6D). Collectively, these data indicate that AAV9-CAG-ROSTASK1 expression in sensory neurons effectively reduces both inflammatory and early neuropathic pain behavior (Figure 6F).

Tissue damage often triggers a heightened pain state in the central nervous system, known as central sensitization (Latremoliere et al, 2009). This amplified pain sensitivity can continue after the initial injury resolves in a phenomenon called latent sensitization, which is due to compensatory inhibitory G protein receptor signaling (Gerum et al, 2022; Nelson et al, 2023). For instance, constitutive activity of mu opioid receptors suppresses spinal nociceptive signaling for months following the resolution of CFA-induced hypersensitivity (Corder et al, 2013). Given the ability of ROSTASK1 to reduce persistent inflammation, we wondered whether it might also prevent the development of latent sensitization. To test this, we administered the opioid receptor inverse agonist naltrexone (3 mg/kg) to mice transfected with either AAV9-CAG-ROSTASK1 or AAV9-CAG-GFP six weeks after CFA injection, following the complete resolution of CFA-induced mechanical hypersensitivity. Naltrexone administration reinstated mechanical hypersensitivity in these mice, but this was markedly reduced in AAV9-CAG-ROSTASK1 mice (Figure 6G,H), suggesting that ROSTASK1 expression in DRG neurons attenuates latent pain sensitization and the propensity for the development of long-term chronic pain.

## Discussion

We have engineered a ROS-activated K^+^ channel, ROSTASK1, that can offset the overproduction of ROS and its pathologic effects on cells. To achieve this functionality, we mutated the parent TASK1 channel to remove its basal activity and tuned the gating mechanism to be sensitive to supraphysiological levels of ROS. Guided by existing structural and functional data, we reinforced the contacts between M2 and M4 to stabilize the X-gate, so that only elevated, and not physiological, concentrations of cellular ROS could open it. We assessed the functionality of our construct by elevating ROS levels using UV-A irradiation, which generates predominantly ^1^O_2_ ROS (Chen et al, 2021) and requires endogenous photosensitizer molecules, such as riboflavin and heme. This process takes several minutes, explaining the slow activation of ROSTASK1 that we observed. Because channel activation could also be achieved using RB – a photosensitizer that generates ^1^O_2_ – and eliminated by ROS scavengers, our results support the hypothesis that ^1^O_2_ activates the channel. The accelerated ROSTASK1 activation we observe upon fusing the channel to miniSOG, which generates a burst of ¹O_2_ under blue light illumination, lends further support to this conclusion. When expressed in mitochondria, ROSTASK1 depolarizes these organelles and reduces excessive ROS production. When expressed at the plasma membrane, the channel hyperpolarizes peripheral neurons and reduces mechanical hypersensitivity associated with inflammatory and neuropathic pain. Thus, ROSTASK1 serves as both a sensor and modulator of ROS levels to mitigate oxidative stress.

### Mechanism of activation

Our data show that ROS activation of ROSTASK1 is irreversible and mediated by two methionine residues (Met156 and Met159) in the vicinity of the X-gate at the interface between the lipid membrane and cytosol. Mutation of these residues to isoleucine or glutamic acid revealed that ^1^O_2_ causes double oxidation of both methionines to sulfones and that another species undergoes oxidation to contribute a small, additional effect. We presume this is why physiological levels of ROS fail to activate ROSTASK1. Moreover, because cellular ^1^O_2_ is short lived (<100 ns), acting locally within cells and tissues (Moan 1990; Gao et al, 2014), significantly elevated levels would be required to achieve double oxidation of methionine residues. This would occur in pathological conditions as well as during experimental manipulation by UV-A irradiation or the combination of visible light and an adequate photosensitizer compound.

These two methionines, Met156 and Met159, form a hydrophobic clamp that stabilizes the opposing Leu241, keeping M3 connected to M4. Both methionines are conserved in TASK3, which shares a high sequence and structural homology with TASK1 and are mutated (M156V and M159I) in patients with *KCNK9* imprinting syndrome (Cousin et al, 2022). Interestingly, TASK3 channel carrying the patient’s variant M156V is WT-like. We speculate that the pathological impact of the mutation may only become apparent during imbalanced regulation of TASK3 by ROS. Indeed, both TASK1 and TASK3 localize to the IMM where they contribute to mitochondrial ROS regulation (Yao et al, 2017), (Yu et al, 2021), and both are known to form heterodimers with intermediate properties (Rinnè et al, 2015), suggesting that the ROS response might be fine-tuned in different (patho)physiological scenarios.

### Powerful optogenetic tool

By adding the ^1^O_2_ generator miniSOG to the ROSTASK1 cytosolic C-terminus, we generated a channel that can be activated by blue light for optogenetic applications. OptoTASK1 has favourable properties for long-lasting neuronal inhibition: low light intensity threshold (50 mW/cm^2^), fast activation (15 sec), irreversible opening (up to 4 hours), and large currents due to robust trafficking to the plasma membrane. Furthermore, canonical intracellular sorting signals were able to direct ROSTASK1 and OptoTASK1 to the IMM, where they were silent at physiological ROS levels, but became active upon UV-A radiation to depolarize the organelle. This permits ROSTASK1 to be used as a smart sensor to both detect elevated levels of ROS and trigger a repair reaction.

At least three other genetically-encoded photosensitizers have been reported – tdKillerRed, SuperNova, and mCherry – all of which generate O2•^−^ and ^1^O_2_ upon green/yellow light exposure with an absorption peak around 590 nm (Onukwufor et al, 2020). Because miniSOG absorbs blue light at 448 nm, with no significant absorption beyond 500 nm, the fusion of tdKillerRed, SuperNova, or mCherry to TASK1 would widen the palette of the synthetic channels for optogenetic applications. This would allow these channels to be activated with longer and more tissue-penetrating wavelengths without the need for chemical photosensitizers. In addition, ROSTASK1 could be activated by multiple combinations of photosensitizer and light beyond the RB/green light and MB/red light that we tested. The possible combinations are as many as the number of photosensitizer compounds, which are largely used in photodynamic therapy (Correia et al, 2021).

### Cellular and whole animal applications

We have demonstrated the effectiveness of ROSTASK1 for reducing ROS production in a cellular model of mitochondrial dysfunction characterized by the m.9176T→C mutation associated with LS. The treatment of mitochondrial diseases is a significant challenge due to the complex nature of their pathophysiology, in which oxidative stress, impaired energy production, and cellular damage intersect. The ability of ROSTASK1 to restore ROS levels in cells harbouring this specific mutation underscores the potential of such a targeted approach in comparison to broad-spectrum antioxidants that have consistently failed to produce substantial clinical benefits in mitochondrial disorders (Grimm et al, 2013; Klopstock et al, 2011). The application of ROSTASK1 could be expanded to a wider spectrum of neurodegenerative diseases in which oxidative stress is a key factor, such as Alzheimer’s and Parkinson’s diseases. Future research could investigate the potential of using ROSTASK1 in combination with other therapeutic strategies, including gene therapy and pharmacological agents that enhance mitochondrial function, to more comprehensively address the multifactorial nature of these conditions.

Our study also highlights the utility of ROSTASK1 as a therapeutic intervention in *in vivo* models of chronic pain. While ROSTASK1 did not affect acute pain behaviors, it robustly reduced the development of neuropathic pain, inflammatory pain, and latent pain sensitization. Notably, the analgesic effect of ROSTASK1 in neuropathic pain was transient, diminishing after 14 days. This finding suggests that ROS play a critical role in the initiation, but not the maintenance, of chronic neuropathic pain. This hypothesis aligns with previous findings that microglia are the predominant source of ROS following peripheral nerve injury and that microglial NADPH oxidase 2 (Nox2)-derived ROS peak immediately after peripheral nerve injury and return to nearly control levels within 14 days (Kim et al, 2010). Furthermore, our results demonstrate that ROSTASK1 effectively prevents chronic inflammation-induced latent sensitization, underscoring its potential to disrupt the transition from acute to chronic pain and reduce the likelihood of relapse in persistent pain conditions. Collectively, these data establish ROSTASK1 as a promising analgesic and a powerful tool for elucidating the mechanisms underlying the initiation and progression of chronic pain at both peripheral and central nervous system levels.

### Future perspectives

ROSTASK1 is a silent channel that can detect pathologic levels of ROS and activate cell- and organelle-specific rescue mechanisms. The fact that ^1^O_2_ has a short lifetime, diffusing only short distances, confines ROSTASK1 activation to specific cellular compartments and/or very small cell volumes. These features guarantee a high degree of cellular or subcellular specificity to this novel ROS-induced ROS control (RIRC) system.

Sonodynamic therapy (SDT) is emerging as a non-invasive cancer treatment with the advantages of deep penetration, good therapeutic efficacy, and minimal damage to normal tissue. This approach utilizes both inorganic and organic sonosensitizers that absorb the energy carried by ultrasonic radiation and release it to generate ROS such as ^1^O_2_, •OH, and •O_2_^−^ (Chen et al., 2023). We envisage that SDT could become an additional application for ROSTASK1, as deep-penetrating ultrasound can be used to potentiate RIRC exerted by the channel.

## Acknowledgements

This work was supported by ERC-2020-PoC n. 966841 and ERC-2023-SyG n. 101118744 to A.M.

## Authors contribution

Conceptualization: A.M.; Methodology: A.S., A.P., A.R., Y.T. and R.K.; Validation: S.T. and A.R.; Investigation: A.R., S.T., T.S.N., H.N.A., C.V. and R.O.; Resources: Y.T. and R.K.; Writing original draft: A.M. and G.T.; Visualization: A.R., A.P.; Supervision: A.M., A.S. and G.T.; Funding acquisition: A.M.

## Declaration of interest

The authors declare no competing interests

## STAR methods

### Chemicals

All chemicals were purchased from Sigma-Aldrich or Fisher Scientific unless otherwise noted. The stock solution of N-acetil-L-cisteine (100 mg/mL) was made by adding ddH2O and stored as aliquots at −20°C. The stock solution of Rose Bengal (1 mM) was made by adding ddH2O and stored as aliquots at −20°C. The stock solution was protected from light and diluted in the bath solution just before the experiments. The stock solution of TROLOX was made by adding DMSO (50 mg/mL). Hydrogen peroxide (30%) was diluted in the bath solution to a final concentration of 10 mM just before experiments and had to be freshly prepared just before experiments. The stock solution of tetramethylrhodamine methyl ester (TMRM, 5 mM) was made by adding DMSO and stored as aliquots at −20°C. Carbonyl cyanide 4-(trifluoromethoxy)phenylhydrazone (FCCP, 10 mM) was made by adding DMSO and stored as aliquots at −20°C. The stock solution (3 mM) of Methylene Blue was made by adding ddH2O and stored as aliquots at −20°C.

### Constructs

The mouse TASK1 (mTASK1) gene was cloned in pcDNA3.1 (for in vitro experiments carried out in HEK293T cells) or in PAAV CAG (for in vivo experiments following viral infection) expression vectors. Site-directed point mutations were introduced using the QuikChange XL Site-Directed Mutagenesis Kit following the specifications recommended by the manufacturer. All constructs were verified by full-length sequencing. Gene cloning, gene truncation and gene fusion with other sequences were carried out by means of Gibson assembly. ROSTASK1 is a truncated version of mTASK1 (mTASK1 1-259) carrying two mutations, that is Q126M and T248F) and with an eGFP fused at its C-terminus. OptoTASK1 bears the same sequence of ROSTASK1 but has miniSOG fused to its C-terminus instead of the eGFP. The mitochondrial version of both ROSTASK1 and OptoTASK1 (mtROSTASK1 and mtOptoTASK1) are characterized by a third mutation, that is H98N, and they are fused to a 6-repeat mitochondrial targeting signal at their N-terminus. Stbl2 competent cells (Invitrogen) were used to amplify plasmid DNA, which was then extracted using Exprep Plasmid SV kit (GeneAll) according to the manufacturers recommended protocol.

### HEK293T Cell culture and transfection

HEK293T cells were cultured in Dulbecco’s modified Eagle’s medium (Euroclone) supplemented with 10% fetal bovine serum (Euroclone), 1% Pen Strep (100 U/mL of penicillin and 100 µg/ml of streptomycin) and stored in a 37°C humidified incubator with 5% CO_2_. Every two- or three-days cells were trypsinized and split to avoid overgrowth. After 20–25 splits cells were discarded, and a new fresh line was thawed to substitute the old one. Cells were grown in a 25 mm^2^ flask and transferred in 35 mm Petri dishes (Sarstedt) the day before transfection. When ~ 70% confluent, HEK293T cells were transfected with wild-type and/or mutant cDNA using Turbofect transfection reagent (Thermo Fisher) according to the manufacturers recommended protocol. For each 35 mm Petri dish 1 µg (for the construct fused to a eGFP) or 1 µg of the gene-containing vector and 0.3 µg of eGFP-containing plasmid (pmaxGFP, AmaxaBiosystems) were used. The constructs fused to miniSOG were co-transfected with a plasmid containing tagRFP to check the efficiency of the transfection and select the cell to record without stimulating the production of ROS by miniSOG. 24–48 hour after the transfection the cells were dispersed by trypsin treatment. Green, fluorescent cells were selected for patch-clamp experiments at room temperature (about 25°C).

### Electrophysiology in HEK cells and data analysis

Currents were recorded in **whole-cell** configuration either with an Axopatch 200B amplifier (Molecular Devices, CA, USA) or with a ePatch amplifier (Elements, Cesena, Italy); data acquired with the Axopatch 200B amplifier were digitized with an Axon Digidata 1550B (Molecular Devices, CA, USA) converter. All data were analysed off-line with Axon pClamp 10.7. Patch pipettes were fabricated from 1.5 mm O.D. and 0.86 I.D. borosilicate glass capillaries (Sutter, Novato, CA, USA) with a P-97 Flaming/Brown Micropipette Puller (Sutter, Novato, CA, USA) and had resistances of 3–6 MΩ. For the experiments in whole-cell configuration the pipettes were filled with a solution containing: 155 mM KCl, 5 mM egtazic acid (EGTA), 3 mM MgCl_2_, 1 mM ATP (Magnesium salt) and 10 mM HEPES–KOH buffer (pH 7.2). The extracellular bath solution contained 150 mM NaCl, 5 mM KCl, 1 mM CaCl_2_, 3 mM MgCl_2_ and 10 mM HEPES–KOH buffer (pH 7.4). The stock solution was prepared solving the powder in milliQ water to obtain a final concentration of 100 mM and adjusting the pH. Single-use aliquots were made and stored at −20°C until the day of the experiment. All measurements were performed at room temperature. For the experiments in **inside-out** configuration the pipette was filled with a solution containing 150 mM NaCl, 5 mM KCl, 1 mM CaCl_2_, 3 mM MgCl_2_ and 10 mM HEPES–KOH buffer (pH 7.4) and the extracellular bath solution contained 155 mM KCl, 5 mM egtazic acid (EGTA), 3 mM MgCl_2_ and 10 mM HEPES–KOH buffer (pH 7.2). For single channel recordings in **cell-attached,** pipettes with a resistance of 12-20 MΩ were used. The pipettes were filled with a solution containing 55 mM NaCl, 100 mM KCl, 1 mM CaCl_2_, 3 mM MgCl_2_ and 10 mM HEPES–KOH buffer (pH 7.4).

**Light treatment** at different wavelengths was performed by means of SPECTRA X light engine (Lumencor) which has different light outputs and enables the precise regulation of light intensity. The light engine was coupled into an optical fiber which was mounted on a manual manipulator, allowing for a precise positioning of the fiber near the recorded cell. Light intensity was measured by means of PM100D Handheld Optical Power Meter (Thorlabs) before the experiments.

The t_1/2_ values were obtained by **fitting** the kinetics of activations with a logistic function**: y= [(A_1_ – A_2_)/1 + (x/x_0_)^p^] + A_2_** where A_1_ is the initial value, A_2_ is the final value, x_0_ is the center and p is the power of the exponential. The data were compared by means of one-way ANOVA followed by Fisher’s test or using Student’s t-test. Significance level was set to p = 0.05. All data are presented as mean ± standard error of the mean (SEM).

### Confocal microscopy

Cell fluorescence was measured using a Nikon Eclipse-Ti inverted confocal microscope interfaced with an A1 series of confocal laser point scanning system for excitation at 405, 488, 561 and 640 nm. Glass bottom Petri dishes containing HEK293T cells were transiently transfected with ROSTASK1, mtROSTASK1 and mtOptoTASK1. Fluorescence analysis was carried out 24 hours after transfection on living cells. The plasma membrane was stained with CellMaskTM Deep Red from Invitrogen according to the manufacturers protocol. Mitochondria were stained with TMRM (15 nM) according to the manufacturers’ protocol. The samples were observed with a 60 × 1.4 NA oil immersion objective (Nikon System). The pinhole aperture was set to 1.0 Airy. The images were collected using low excitation power (488 and 640 nm) at the sample and acquiring the emission range through bandpass filters 525/50 and 700/75 for EGFP and CellMaskTM emission, respectively, by means of built-in GaAsP PMT detectors of the confocal microscope. For the experiment of photoactivation of mtROSTASK1 with UV-a light we used a photoactivation protocol in which the sample was continuously illuminated with a 405 nm laser (30 mW cm^−2^) and every 2 minutes the image was acquired. For the analysis, the variation of the TMRM fluorescence of a single cell expressing mtROSTASK1 was normalized and expressed as a ratio between the fluorescence of a cell expressing mtROSTASK1 and a nearby cell not expressing the channel. The average of the ratios was then calculated. As a control the same analysis was carried out by comparing two nearby cells not expressing the channel using the same protocol as described above.

### Cell Lines

Human healthy and m.9176T→C myoblasts were obtained from muscle biopsies conducted during diagnostic path at the Fondazione IRCCS Ca’ Granda Ospedale Maggiore Policlinico in Milan, Italy. Healthy muscle biopsies were collected from individuals undergoing orthopedic surgery, and m.9176T→C myoblasts were obtained from a previous study (Ronchi et al, 2011).

The muscle tissue samples were processed in Dulbecco’s modified Eagle’s medium high glucose (DMEM) under sterile conditions. After removal of fat cells and fibrotic tissue, the muscle tissue was mechanically minced, digested in 1 mg/ml collagenase I (Sigma-Aldrich) in Dulbecco’s Modified Eagle Medium (DMEM) with high glucose, 10% FBS, and 1% Penicillin/Streptomycin at 37°C for 70 min. The resulting digests were washed with PBS, and the suspensions were filtered through a 40 µm nylon mesh. After lysing erythrocytes with ACK lysing buffer (ThermoFisher) for 3–5 min on ice and washing with PBS, myoblasts were isolated from other cells using flow cytometry cell sorting, identified as cells expressing CD56+, CD146+, and CD82+. Cells were cultured in F10 supplemented with 10% FBS, 1% Pen Strep (Gibco USA), and 5 ng/mL EGF (Sigma-Aldrich).

### Cell Lines Transfection

Healthy and m.9176T→C myoblasts were seeded at a density of 10,000 cells per well in a 96-well multiwell plate and transfected with GFP or Mito-ROS-Task DNA using Lipofectamine 2000 (ThermoFisher) after 24 h. A mix of 50 ng of DNA and 0.2 μl of Lipofectamine 2000 in OptiMem medium was incubated for 20 min at room temperature. The resulting DNA/Lipofectamine solution (10 μl) was directly added to each well. ROS levels were evaluated 72 h post-transfection.

### ROS Assay

ROS levels in healthy and m.9176T→C myoblasts, 72 h post-transfection with GFP or Mito-ROS-TASK, were assessed using the ROS-GloTM H2O2 Assay (Promega) following a non-lytic procedure. Media samples were transferred to a separate white plate after 4 h of incubation, and relative luminescence units were measured by a GloMax Discover plate reader.

### MTT Assay

The proliferative capacity of cells was assessed using MTT experiments. Briefly, 50 μg/mL MTT solution (Methylthiazolyldiphenyl-tetrazolium bromide) (Sigma Aldrich) was added to healthy and m.9176T→C myoblasts in a 96-well plate and incubated for 4 hours at 37 °C. The dye was solubilized with DMSO (Sigma Aldrich), and absorbance at 560 nm was recorded with a GloMax Discover plate reader. The MTT assay was performed after the ROS-GloTM H2O2 Assay following a non-lytic procedure.

### Animals

Adult C57Bl/6J (Jackson Laboratory, # 000664) male and female mice were group housed, provided access to food and water ad libitum, and maintained on a 12:12 hour light:dark cycle in temperature and humidity controlled rooms. Equal numbers of male and female mice were used in all behavioral experiments. Although we were not powered to detect significant sex differences, no major/obvious trends in sex differences were observed and means from both sexes were pooled. All procedures were consistent with the guidelines for the treatment of animals of the International Association for the Study of Pain and were approved by the University of Florida Institutional Animal Care and Use Committee.

### Intrathecal Adeno-associated Virus (AAV) Administration

Purified adeno-associated virus (AAV) packaging for in vivo testing was performed by Charles River. Packaged AAV9-CAG-ROSTASK1 arrived at a virus titer of 1.032 x 1013 GC/mL and was used at this titer for experiments. AAV9-CAG-GFP arrived at a virus titer of 5.64 x 1013 GC/mL and was diluted 1:5 in sterile PBS for experimental testing. Viruses were stored at −80°C until injection. Intrathecal injections were performed in lightly restrained unanesthetized mice. Briefly, a 30G needle attached to a Hamilton microsyringe was inserted between the L5/L6 vertebrae at the cauda equina, puncturing the dura (confirmed by presence of reflexive tail flick). We then injected a 5 ml volume of AAV9 before returning the mice to their home cage. Behavioral testing was performed 21 days post-intrathecal injections to permit stable transfection.

### Pain Injury Induction

Spared Nerve Injury: SNI was performed as previously described (Nelson et al, 2023; Qi et al, 2023). Briefly, mice were anesthetized with inhaled isoflurane (5% induction and 2% maintenance) and the left hind limb was shaved with trimmers and asepticized with 70% ethanol and a ChloraPrepTM swabstick (BD Cat: 260100). A small incision was made in the skin of the hind left leg and the underlying muscle was spread via blunt dissection to expose the underlying branches of the sciatic nerve. The peroneal and tibial nerves were then ligated with 6-0 silk sutures (Butler Schein Animal Health Cat:NC0049524) and transected while carefully avoiding the sural nerve. The muscle tissue was then loosely sutured with 5-0 vicryl sutures (Med Vet International Cat:50-118-0847) and the skin was closed with 9 mm wound clips (Braintree Scientific Cat:NC9281117). Topical triple antibiotic ointment (Neosporin Neomycin Sulfate/Bacitracin Zinc/Polymycin Ointment; Hanna Pharmaceutical Supply Co. Cat:NC0100117) was applied to the wound. Wound clips were removed 7-10 days post-surgery and behavioral experiments began 3 days after surgery.

Complete Freund’s Adjuvant (CFA) Administration: Briefly, mice were anesthetized with inhaled isoflurane (5% induction and 2% maintenance) and injected subcutaneously with 20 mL of undiluted Freund’s Complete Adjuvant (MilliporeSigma Cat:AR001) into the midplantar region of the left hindpaw with a 30 G needle attached to a 25 mL Hamilton syringe. Behavioral experiments began 3 days after surgery.

Sham Injury: Mice were anesthetized with inhaled isoflurane (5% induction and 2% maintenance) for 5 minutes and then returned to their home cage. Behavioral experiments began 3 days after surgery.

### Behavioral Testing

Mechanical Withdrawal Threshold: Testing was performed as previously described (Nelson et al, 2022; Nelson et al, 2023; Fu et al 2019). Mice were habituated to plexiglass chambers (80 × 80 × 110 mm) on a raised wire mesh platform for 60 minutes immediately prior to behavioral testing. Testing was performed using a calibrated set of logarithmically increasing von Frey monofilaments (Braintree Scientific Cat: 58011) that range in gram force from 0.007 to 6.0 g. Beginning with a 0.4 g filament, these were applied perpendicular to the lateral hindpaw surface with sufficient force to cause a slight bending of the filament. A positive response was denoted as a rapid withdrawal of the paw within 4 seconds of application and was followed by application of the next lower filament. A negative response was followed by application of the next higher filament. An up-down method (Chaplan et al, 1994) was used to calculate the 50% withdrawal threshold for each mouse.

Heat Withdrawal Threshold: We evaluated acute nociceptive responses with a hot plate as previously described (Nelson et al, 2022). Briefly, prior to testing mice acclimated to the testing room for at least 30 minutes to minimize stress-induced behaviors. Following acclimation, mice were gently placed on a 52°C Ugo Basile hot/cold plate (Stoelting Cat:55075). The trial ended when the mouse exhibited a nociceptive response (such as paw licking or jumping) or after a cutoff time of 30 seconds if no response was observed. Testing was repeated for 3 trials per mouse with intervals of at least 10 minutes between each trial. The latency to respond for each trial was averaged to produce the mean latency for statistical analysis.

Motor Coordination Testing: We assessed motor coordination using a Ugo Basile accelerating Rota-Rod (Stoelting Cat: 57624) as previously described (Nelson et al, 2022). Briefly, mice were introduced to the apparatus by placing them on the wheel set to an initial speed of 4 RPM for 30 seconds. After this brief acclimation period, the rotarod initiated acceleration, gradually ramping up over 300 seconds to a final speed of 40 RPM. Each trial concluded upon the mouse’s displacement from the rotating wheel. Five successive trials were conducted per mouse.

Latent Sensitization: To assess for chronic latent hyperalgesia, we subcutaneously administered an opioid inverse agonist (naltrexone hydrochloride dissolved in sterile saline; 3 mg/kg; Thermo Scientific Chemicals Cat:J60590-MD) after the resolution of CFA-induced hypersensitivity (6 weeks post-CFA) as previously described (Walwyn et al, 2016). Subsequently, mechanical withdrawal thresholds were assessed as described above.

## Notes

### Competing Interest Statement

The authors have declared no competing interest.

