## Supplementary material for "A synthetic potassium channel reduces oxidative stress via cellular adaptronics": Supplmental information

### Supplemental information

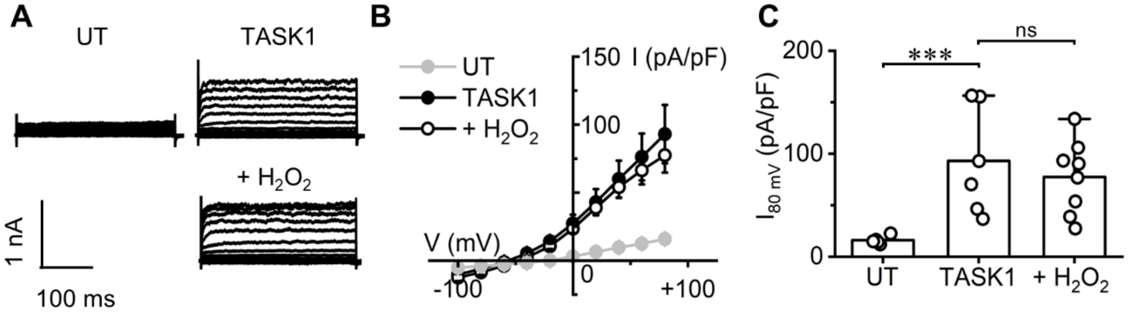


**Figure S1 - TASK1 channel is insensitive to 10 mM $H_2O_2$**

**A**: Representative current traces of HEK293T cell expressing TASK1 in control solution and in the presence of 10 mM $H_2O_2$ (n= 8) in the extracellular solution. An untransfected (UT) cell is shown for comparison. **B**: I/V relationships of experiments in panel A. UT, n= 5; TASK1, n=6; TASK1 + $H_2O_2$, n=8. Data are shown as mean ± SEM. **C**: Current density values measured at + 80 mV from the same data shown in B. Data are shown as mean ± SEM along with single experiments values. One-way Anova with Fisher's LSD test (***$p<0.001$; ns, $p>0.05$).

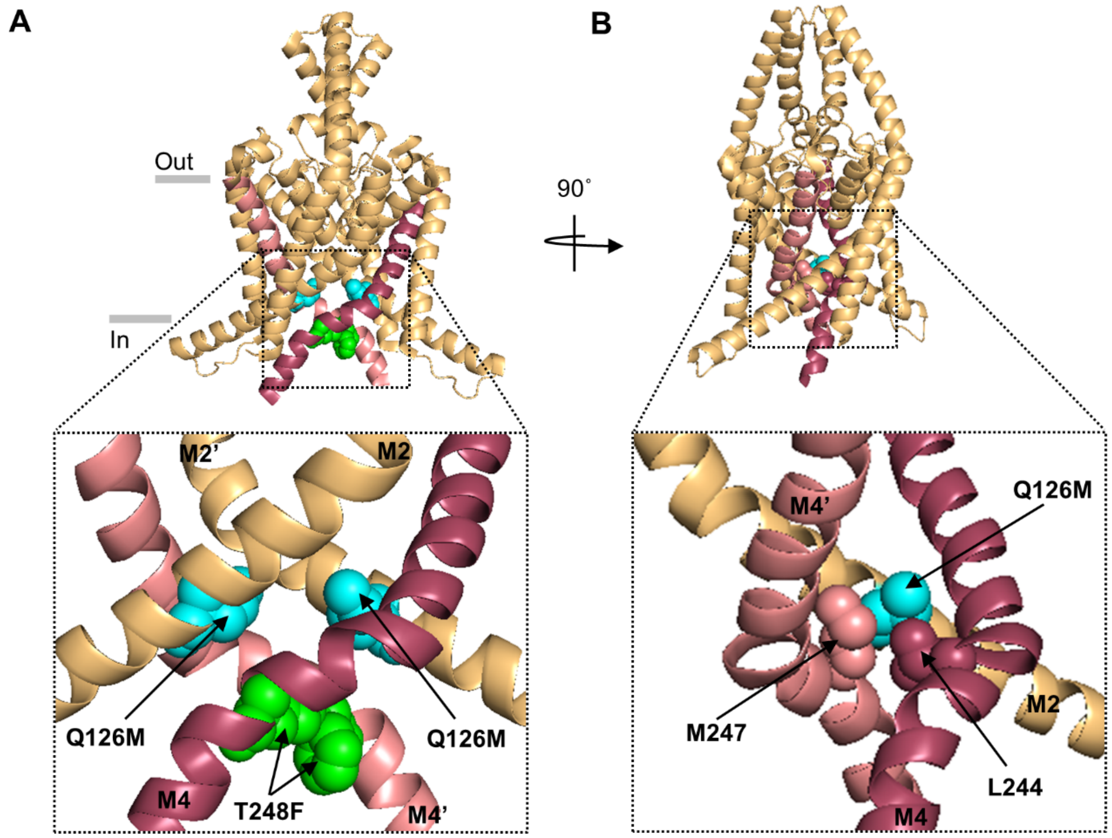


**Figure S2- hydrophobic reinforcement of the X-gate**

**A**: Front view the human TASK1 structure truncated at Glu259 (PDB: 6rv2). M4 and M4' helices are shown in red and pink, respectively, to highlight the X-gate structure. The substitutions Q126M and T248F introduced in ROSTASK1 were manually introduced in the structure using the "mutagenesis" tools of Pymol software. Their side chains are shown, as spheres, in light blue and green, respectively. Boxed, magnification of the X-gate showing that the distance between the two F248 residues may be compatible with strong hydrophobic interactions that stabilize the closed conformation of the X-gate. **B**: 90° rotated view of the structure shown in A. Boxed, magnification showing only M2, M4 and M4' helices. Side chains of M126 on M2, M247 on M4' and L244 on M4 residues are shown as spheres to describe the possible newly introduced hydrophobic interaction leading to the stabilization of the X-gate in the closed conformation.

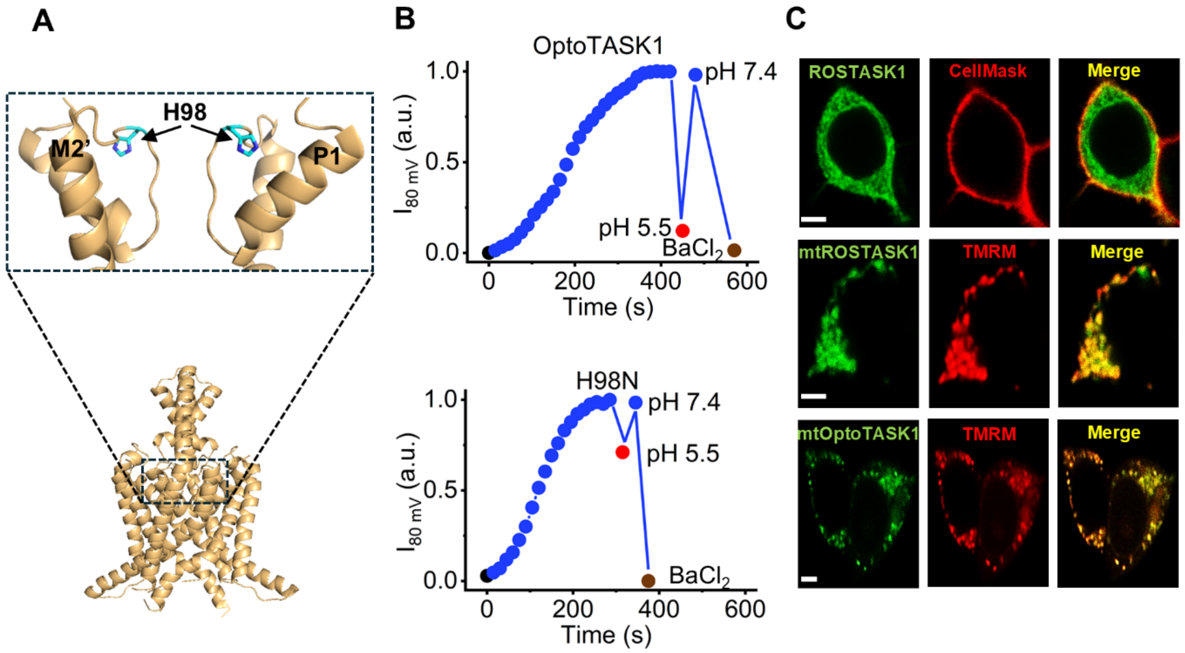


**Figure S3- Removal of extracellular pH dependency from ROSTASK1 and OptoTASK1**

**A:** Structure of human TASK1 channel (PDB ID: 6RV2). Boxed, magnified view of the selectivity filter with side chains of H98 residues are highlighted in blue. For clarity, only M2' and P1 helices are labeled. **B,** effect of low extracellular pH on OptoTASK1 current. Normalized current measured at 80 mV from a representative cell expressing OptoTASK1 wt (top panel) and OptoTASK1 H98N mutant (bottom). Black dot at t0 shows current measured in safe light. Current onset was triggered by 470 nm (55 mW/cm$^2$) light illumination (blue symbols) during perfusion of extracellular solution at pH 7.4. Red symbol: current measured by perfusing extracellular solution at pH 5.5. 10 mM $BaCl_2$ addition abolished OptoTASK1 current (brown dot). **C**: Confocal fluorescence images showing the mitochondrial localization of mtROSTASK1. First row, plasma membrane colocalization of the control channel ROSTASK1 (green) with the red signal of the plasma membrane indicator CellMASK. Middle row, mitochondrial colocalization of mtROSTASK1 (green) with the red signal of the mitochondrial marker TMRM. Third row, same as middle row for mtOptoTASK1. Scale bar: 5 um

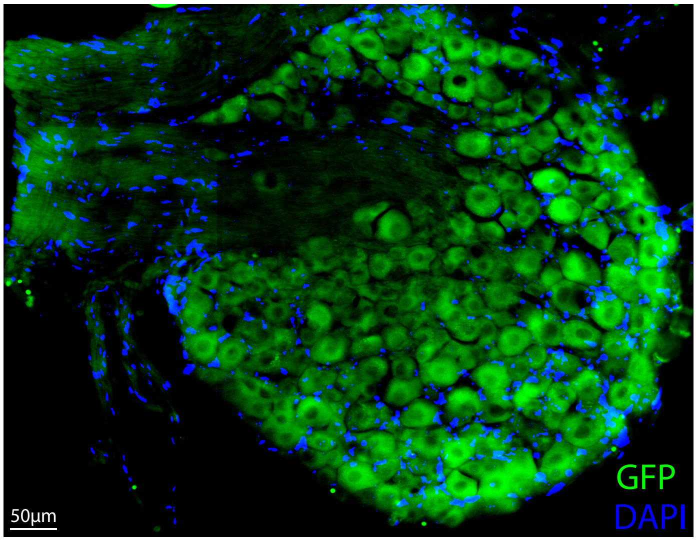


**Figure S4- GFP signal labeled in the dorsal root ganglia.**
Green labeled cells represent virally infected cells, blue represents DAPI stained nuclei.

| | Light intensity mW/cm² | | | | | | | | | | | | | |
|---|---|---|---|---|---|---|---|---|---|---|---|---|---|---|
| | 1 | | 3 | | 10 | | 15 | | 20 | | 30 | | 55 | |
| | $t_{1/2}$ (s) | n | $t_{1/2}$ (s) | n | $t_{1/2}$ (s) | n | $t_{1/2}$ (s) | n | $t_{1/2}$ (s) | n | $t_{1/2}$ (s) | n | $t_{1/2}$ (s) | n |
| TASK1 (395 nm) | - | - | - | - | - | - | - | - | - | - | - | - | 336 ± 20 | 4 |
| TASK1 + RB (555 nm) | - | - | - | - | - | - | - | - | - | - | - | - | 16 ± 3.6 | 7 |
| ROSTASK1 (395 nm) | - | - | - | - | - | - | - | - | - | - | - | - | 431 ± 88 | 6 |
| ROSTASK1 + RB (555 nm) | - | - | - | - | - | - | - | - | - | - | - | - | 25 ± 5.4 | 5 |
| OptoTASK1 (470 nm) | - | - | 822 ± 72 | 5 | 492 ± 48 | 8 | 390 ± 24 | 10 | 348 ± 48 | 4 | 156 ± 18 | 6 | 108 ± 12 | 8 |
| Q103V (470 nm) | 450 ± 72 | 5 | 348 ± 48 | 4 | 174 ± 24 | 9 | - | - | 58 ± 18 | 5 | - | - | 15 ± 4.2 | 3 |
| Q103V W81A (470 nm) | 534 ± 84 | 5 | 276 ± 36 | 4 | 144 ± 18 | 5 | - | - | 60 ± 24 | 3 | - | - | 21 ± 3.6 | 4 |

**Table S1- Half time for ROS-induced activation of TASK1 and TASK1-derived synthetic channels**

Light wavelengths are indicated in nm and provided at the indicated mW/cm$^2$; RB, Rose Bengal was supplied at 100 nM; n, indicates number of repetition (cells)
